# Frameshift Mutations in the Microevolution and Macroevolution of Viruses

**DOI:** 10.1101/2022.07.20.500745

**Authors:** Yu-Fei Ji, Jian-Wei Shao, Rui-Xu Chen, Huan-Yu Gong, Ming-Hui Sun, Guo-Hui Li, Ji-Ming Chen

## Abstract

Some nucleotide insertions or deletions (indels) in open reading frames of virus genes lead to frameshift mutations (FSMs), which can drastically change amino acid sequences. Most FSMs are deleterious and inhibited by natural selection, but some FSMs could aid viruses in adapting to new hosts, escaping immunity, or changing viral pathogenicity or transmissibility. Surprisingly, various fundamental aspects of FSMs in virus evolution remain unknown. In this report, we identified 679 FSMs in the genomes of viruses from 13 randomly selected animal virus families using randomly selected sequences. The 679 FSMs were formed with 1−5 indels, and most (89.4%) FSMs were fixed through the compensatory mechanism with two indels. Each FSM changed 3−209 (mean=15.4, median=13) amino acid residues. FSM frequencies were usually higher in viral sequences with lower sequence identities and steeply increased when sequence identities declined to 60.0%−69.9% or 70.0%−79.9%. This suggests that FSMs are more important to the macroevolution (i.e., inter-species evolution, including speciation) than to the microevolution (i.e., intra-species evolution) of viruses. This study provided novel evidence for the hopeful monster hypothesis in evolutionary biology. Furthermore, we found that FSMs occurred at different frequencies among genes in the same virus genomes and among virus families. Collectively, this study revealed multiple fundamental features of FSMs in virus evolution for the first time and provided novel insights into the mechanisms of macroevolution and speciation.

---

Viruses are the etiological agents of many infectious diseases of humans, animals, and plants, such as rabies, influenza, polio, dengue fever, and coronavirus disease 2019 (COVID-19). Virus evolution involves nucleotide substitution, insertion, and deletion, besides genomic recombination, re-assortment, and rearrangement (1). Insertions or deletions (indels) of 3n±1 nucleotides in open reading frames (ORFs) of protein-coding genes lead to frameshift mutations (FSMs) (2, 3).

As FSMs can drastically change amino acid sequences, most FSMs in virus genomes could be deleterious and inhibited by natural selection. However, some FSMs could aid viruses in adapting to new hosts, escaping immunity, or changing their pathogenicity or transmissibility. For example, FSMs in the genome of vaccinia virus could rapidly attenuate the virus (4). FSMs in the ORF7a gene of severe acute respiratory syndrome coronavirus 2 (SARS-CoV-2) could reduce the virus pathogenicity (5). An FSM in the nsp6 gene of SARS-CoV-2 could be associated with adaptation of the virus to cats, dogs, and hamsters (6). Two FSMs in the nsp2 gene of SARS-CoV-2 compared with SARS-CoV-1 could be associated with the virus transmissibility (7). An FSM in the precore region of the genome of hepatitis B virus could be associated with spontaneous reactivation of the virus (8).

Few studies on FSMs in virus evolution have been reported. A recent milestone study parsed 1,233,275 viral sequences of 10,115 genes of many RNA viruses, and found that FSMs sporadically occurred in 385 genes among the sequences with identities of >97% (9). For instance, two of 5,606 hemagglutinin gene sequences with identities of >97% of H1 subtype influenza virus carried FSMs, and six of 685 the hemagglutinin neuraminidase (HN) gene sequences with identities >97% of paramyxoviruses carried FSMs (9).

Although FSMs are well known to biomedical researchers, various fundamental aspects of FSMs, including their roles in virus evolution, remain unclear. In this report, we investigated some fundamental features of FSMs in virus genomes on four scales, which pertains to the microevolution (i.e., intra-species evolution) and macroevolution (i.e., inter-species evolution, including speciation) of viruses.

## RESULTS

### FSMs in 13 virus families

We downloaded and analyzed randomly selected gene sequences of randomly selected viruses from 11 randomly selected RNA virus families and 2 randomly selected DNA virus families, including the newly established virus families of *Hantaviridae, Nairoviridae, Peribunyaviridae*, and *Phenuiviridae* (10-22). These sequences constituted 604 fas files, which harbored 400, 167, 195, and 244 groups with intra-group sequence identities of 90.0%−99.9%, 80.0%−89.9%, 70.0%−79.9%, and 60.0%−69.9%, respectively (Fig. 1). FSMs occurred in all the 13 virus families, and 679 FSMs were identified in the total of 1006 sequence groups (Fig. 1, Fig. 2A, Table 1). Each of the 679 FSMs changed 3−209 (mean=15.4, median=13) amino acid residues (Fig. 2B). Details of the fas files, sequence groups, and FSMs were shown in Table S1.

**Table 1.** Frequencies of frameshift mutations (FSMs) in viral sequences with different sequence identities^a^

| Virus family | Gene | Sequence length | Sequence used | FSM counts | FSM frequencies as per sequence identities |  |  |  |
| --- | --- | --- | --- | --- | --- | --- | --- | --- |
|  |  |  |  |  | 90.0%–99.9% | 80.0%–89.9% | 70.0%–79.9% | 60.0%–69.9% |
| *Arenaviridae* | Large protein | 6616 | 167 | 80 | 0.11 (6/42) | 0.27 (5/14) | 0.14 (4/22) | 1.96 (65/25) |
| *Circoviridae* | Capsid protein | 752 | 130 | 11 | 0.00 (0/28) | 0.00 (0/11) | 2.66 (4/10) | 3.58 (7/13) |
| *Coronaviridae* | Spike protein | 3845 | 169 | 70 | 0.18 (5/37) | 0.39 (3/10) | 0.91 (7/10) | 3.58 (55/20) |
| *Flaviviridae* | Polyprotein | 10120 | 172 | 49 | 0.07 (5/36) | 0.16 (4/12) | 0.10 (2/10) | 0.89 (38/21) |
| *Hantaviridae* | Large protein | 6455 | 85 | 31 | 0.09 (2/17) | 0.15 (2/10) | 0.22 (4/14) | 1.27 (23/14) |
| *Nairoviridae* | Large protein | 11840 | 77 | 94 | 0.00 (0/16) | 0.00 (0/10) | 0.45 (14/13) | 1.88 (80/18) |
| *Orthomyxoviridae* | Hemagglutinin | 1698 | 205 | 2 | 0.00 (0/42) | 0.00 (0/15) | 0.00 (0/11) | 0.33 (2/18) |
| *Paramyxoviridae* | Large protein | 6696 | 224 | 49 | 0.03 (2/50) | 0.19 (3/12) | 0.20 (4/15) | 1.32 (40/22) |
| *Peribunyaviridae* | Large protein | 6761 | 136 | 44 | 0.07 (3/32) | 0.06 (1/13) | 0.06 (2/25) | 1.22 (38/24) |
| *Phenuiviridae* | Large protein | 6266 | 95 | 15 | 0.00 (0/19) | 0.00 (0/12) | 0.00 (0/10) | 1.20 (15/10) |
| *Picornaviridae* | Polyprotein | 6720 | 207 | 126 | 0.07 (3/31) | 0.30 (10/25) | 0.61 (27/33) | 1.94 (86/33) |
| *Poxviridae* | DNA polymerase | 3014 | 66 | 7 | 0.00 (0/22) | 0.00 (0/12) | 0.30 (2/11) | 0.92 (5/9) |
| *Togaviridae* | NS protein | 7425 | 126 | 101 | 0.07 (3/28) | 0.06 (1/11) | 1.53 (25/11) | 2.85 (72/17) |

<sup>a</sup>The data in the parentheses after the frequencies showed the relevant FSM counts versus the relevant group counts.

**FIG 1.**
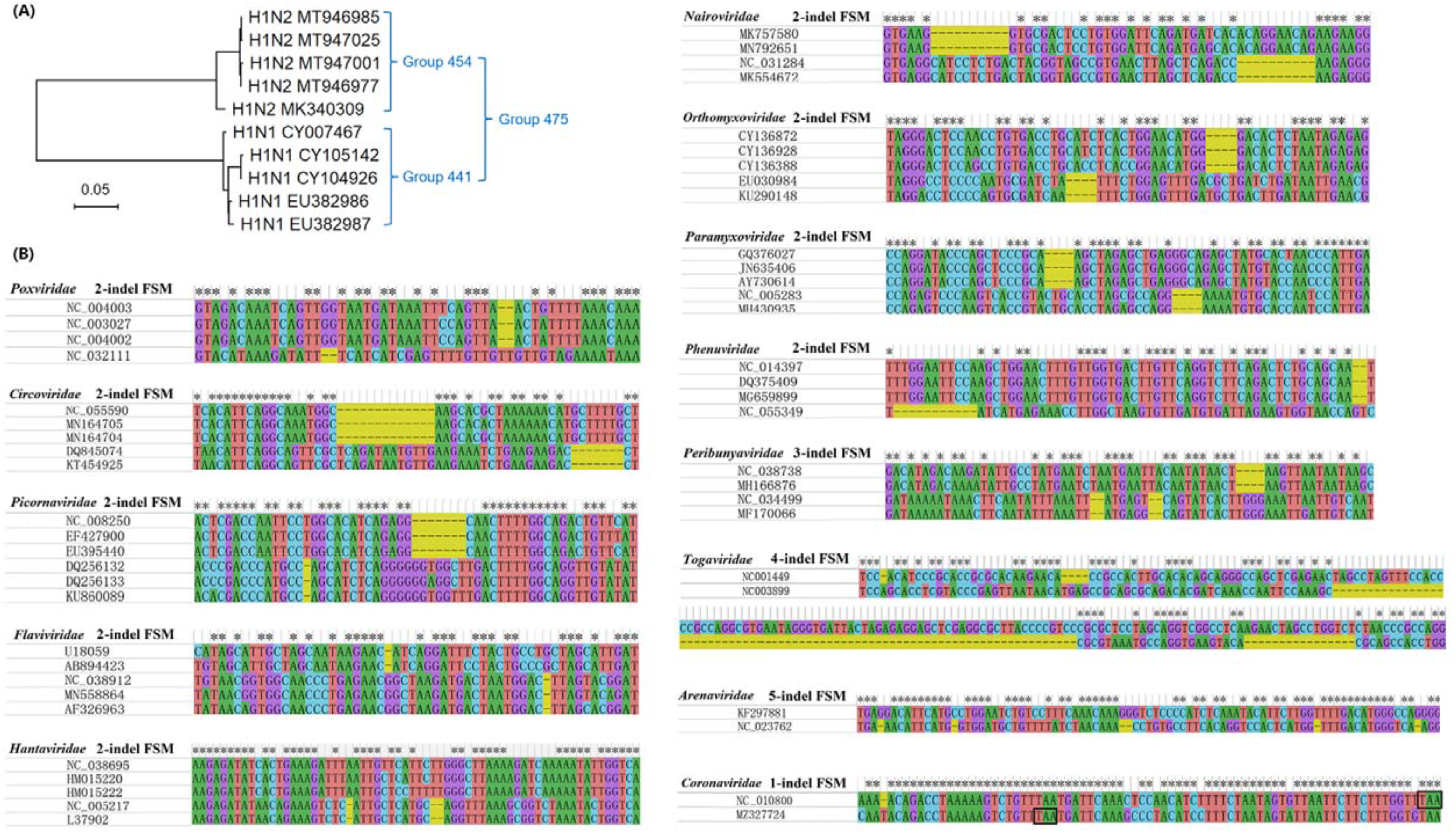
Phylogenetic relationships of three groups harbored in a fas file (A) and examples of frameshift mutations and frameshift types in 13 virus families(B).

**FIG 2.**
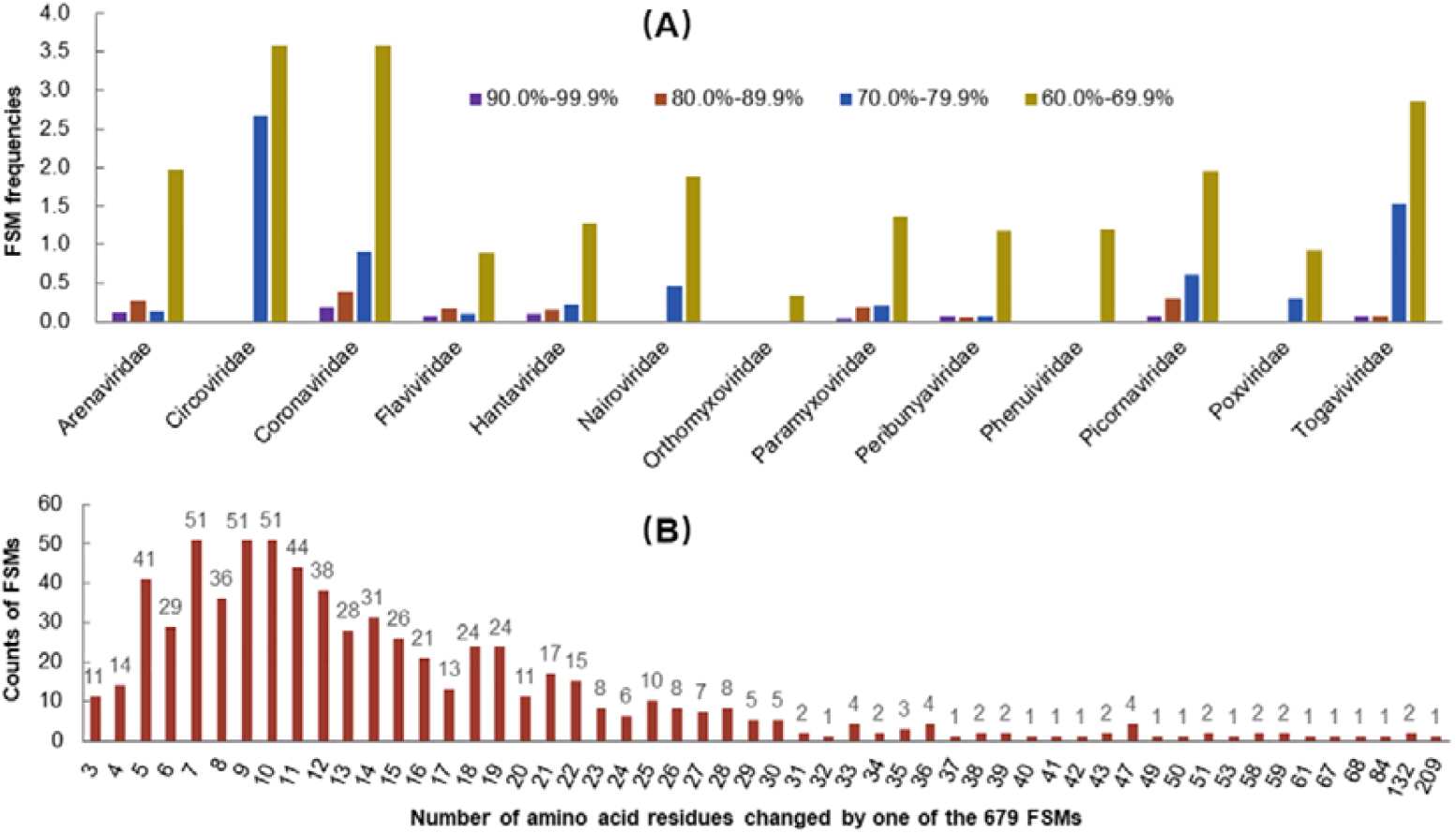
Frequencies of frameshift mutations in 13 virus families among the sequences with four ranges of sequence identities (A) and distribution of the numbers of amino acid residues changed by the 679 frameshift mutations identified in this study (B).

### FSM frequencies and sequence identities

Fig. 2 and Table 1 showed that FSMs usually occurred more frequently in the sequences of the same viral gene with lower sequence identities (P<0.05, by the paired samples Wilcoxon test). For example, FSMs occurred 18.9, 8.2, and 2.9 times more frequently in the spike gene of coronaviruses with sequence identities of 60.0%−69.9% than in the same gene with sequence identities of 90.0%−99.9%, 80.0%−89.9%, and 70.0%−79.9%, respectively. FSMs occurred 26.7, 5.9, and 2.2 times more frequently in the polyprotein gene of picornaviruses with sequence identities of 60.0%−69.9% than in the same gene with sequence identities of 90.0%−99.9%, 80.0%−89.9%, and 70.0%−79.9%, respectively. Additionally, possibly due to the sampling error, FSMs did not always occur more frequently in virus genomes of the same family with lower sequence identities. For example, FSMs occurred less frequently in the groups with sequence identities of 70.0%−79.9% than in the groups with sequence identities of 80.0%−89.9% in *Arenaviridae* (Table 1). Moreover, FSM frequencies increased not gradually with the decline of sequence identities, but steeply when sequence identities declined to 60.0%−69.9% or 70.0%−79.9% (Fig. 2).

### Distribution of FSM types

Of the 679 FSMs, four (0.6%) were 1-indel FSMs, which were formed with a single indel near to termination codons (Fig. 1B). The remaining 675 FSMs were compensatory, and 607 (89.4%) were 2-indel FSMs, 62 (9.1%) 3-indel FSMs, five (0.7%) 4-indel FSMs, and one (0.1%) 5-indel FSMs (Fig. 3). From the percentages of each FSM type in the 13 virus families observed in this study (Text S1), 2-indel FSMs were significantly more prevalent than other types of FSMs, and 3-indel FSMs were significantly more prevalent than other types of FSMs except 2-indel FSMs (P<0.01, by the paired samples Wilcoxon test).

**FIG 3.**
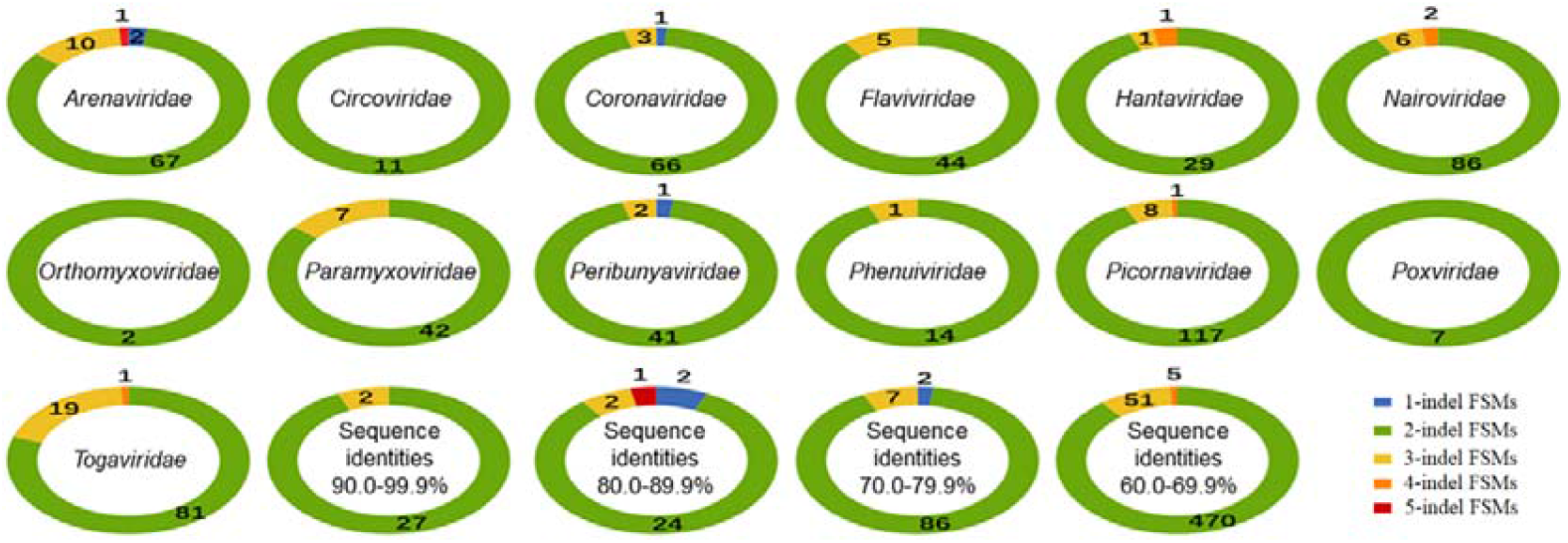
Examples of five types of frameshift mutations (A) and the distribution of frameshift mutations on the four scales of sequence identities (B). Termination codons were marked with rectangles.

### FSM frequencies and virus families

FSMs occurred with different frequencies among virus families. For example, Table 1 showed that FSMs occurred more frequently in the spike gene of coronaviruses than in the hemagglutinin gene of influenza viruses by more than 10 times in their groups with sequence identities 60.0%−69.9% (P=0.001, by the common Wilcoxon test, Text S1). These two genes encode the viral surface proteins to bind to cell receptors. They both evolve more rapidly than most other genes in the viral genomes (23). Similarly, FSMs occurred more frequently in the polyprotein genes of picornaviriuses than in the polyprotein genes of flaviviruses by more than 100% when the sequence identities were 60.0%−69.9% (P=0.007, by the common Wilcoxon test, Text S1). These two polyprotein genes encode all proteins of the relevant viruses (21).

### FSM frequencies and virus genes in the same genomes

We analyzed the FSM frequencies in a viral polymerase gene and a surface protein gene of the above 13 randomly selected families using 39 pairs of randomly selected sequences with identities 60.0%−79.9%. Table 2 showed that FSMs were significantly less frequent in the polymerase genes than in the surface protein genes of the same genomes (P=0.000, by the paired samples Wilcoxon test, Text S1), for most (28/39) of the pairs.

**Table 2.** FSM frequencies of 39 randomly selected pairs of the polymerase gene (PG) and a surface protein gene (SPG) of 13 virus families

| Virus family | Pair 1 |  | Pair 2 |  | Pair 3 |  |
| --- | --- | --- | --- | --- | --- | --- |
|  | PG | SFG | PG | SFG | PG | SFG |
| *Arenaviridae* | 1.7 | 13.9 | 2.3 | 0.0 | 1.5 | 0.0 |
| *Paramyxoviridae* | 1.5 | 6.1 | 3.0 | 3.0 | 0.7 | 2.5 |
| *Hantaviridae* | 0.8 | 7.3 | 2.3 | 10.2 | 0.8 | 7.3 |
| *Nairoviridae* | 0.8 | 8.0 | 1.7 | 6.0 | 0.4 | 7.6 |
| *Phenuviridae* | 1.6 | 5.1 | 0.8 | 0.0 | 2.4 | 9.7 |
| *Peribunyaviridae* | 0.7 | 0.0 | 0.7 | 9.4 | 0.8 | 5.9 |
| *Orthomyxoviridae* | 0.0 | 0.0 | 0.0 | 2.9 | 0.0 | 5.9 |
| *Circoviridae* | 0.0 | 6.9 | 5.6 | 6.9 | 0.0 | 7.2 |
| *Coronaviridae* | 0.0 | 4.0 | 1.1 | 2.5 | 0.8 | 3.8 |
| *Togaviridae* | 2.1 | 2.7 | 2.0 | 5.4 | 0.7 | 4.0 |
| *Poxviridae* | 1.7 | 9.0 | 0.0 | 0.0 | 1.7 | 8.9 |
| *Picornaviridae* | 0.0 | 9.8 | 0.0 | 6.1 | 0.0 | 15.9 |
| *Flaviviridae* | 1.9 | 0.0 | 1.8 | 3.3 | 0.0 | 3.6 |

### FSM identification and sequence alignment

We analyzed 13 randomly selected pairs of sequences with identities 60.0%−79.9% from the above 13 virus families. The results showed that fewer FSMs were identified if viral sequences were aligned using MUSLE with higher gap penalties (P<0.05, by the paired samples Wilcoxon test, Table 3, Text S1), likely because higher gap penalties can lead to the identification of fewer indels. Table 3 also showed that the numbers of FSMs identified using MAFFT with the mode of E-INS-I were highly consistent with the FSMs identified using MUSCLE with the gap penalty of −800, and fewer than the FSMs identified using MAFFT with the mode of FFT-NS-2 (P<0.05, by the paired samples Wilcoxon test, Text S1).

**Table 3.**
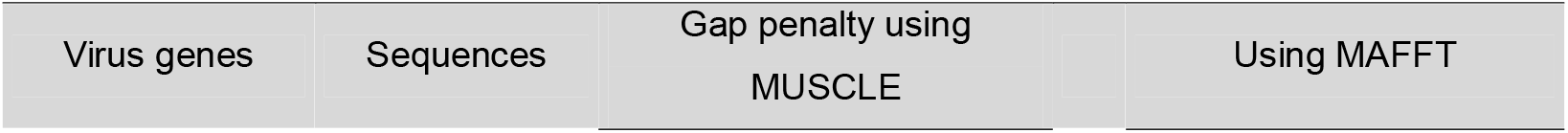
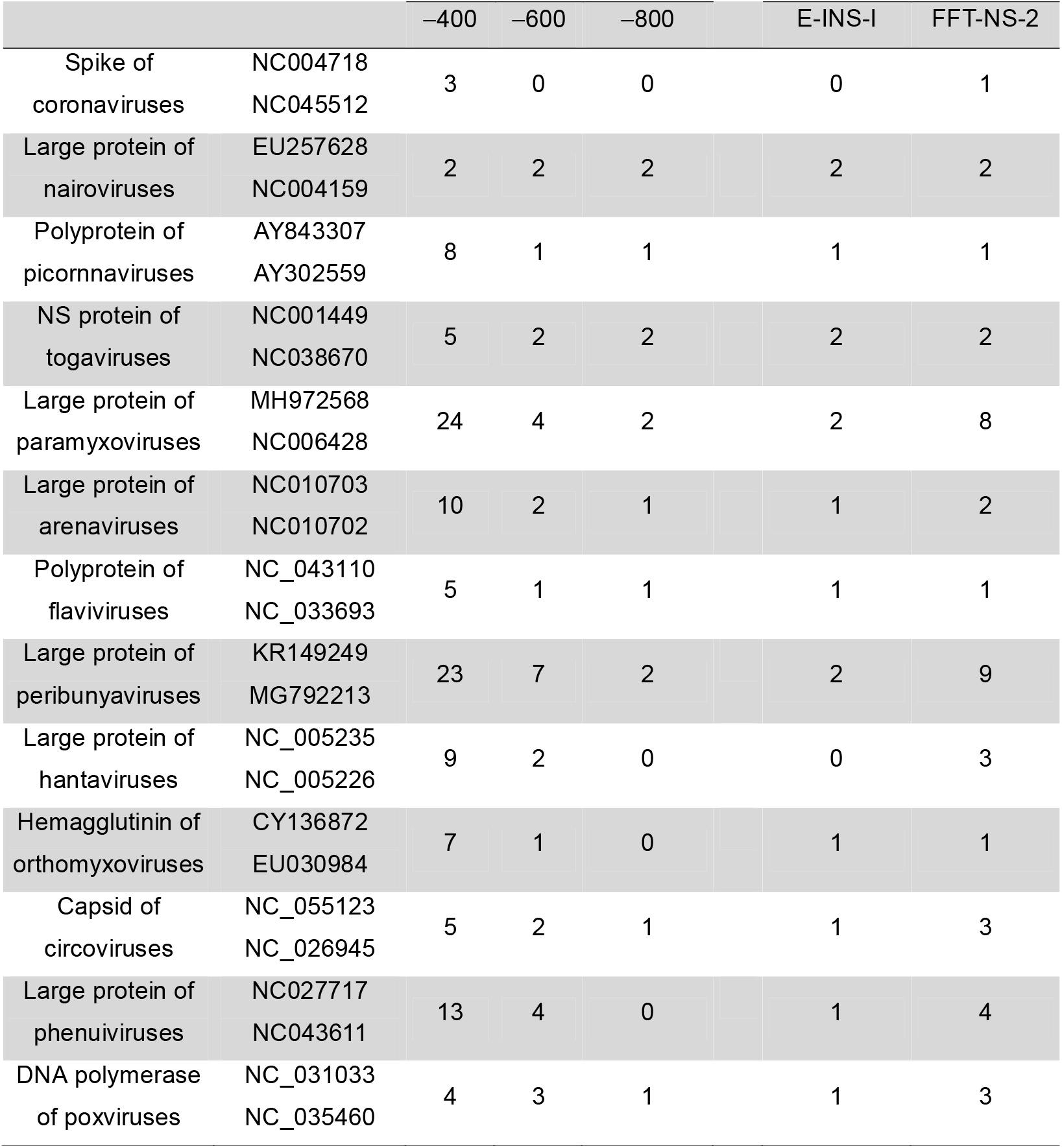
Counts of FSMs in viral sequences aligned using two software tools

## DISSCUSSION

Few studies have been reported regarding FSM frequencies in the genomes of viruses or cellular organisms, although the potential significance of some FSMs of viruses or cellular organisms has been investigated. (1-8). This could be partially because researchers usually focus on the microevolution of viruses or cellular organisms (e.g., microevolution of influenza viruses or SARS-CoV-2), and FSMs were relatively rare in microevolution, as revealed by this report and a previous report (9). Consequently, this study could reveal various fundamental features of FSMs in virus evolution for the first time from a macroevolution or broader view.

A previous report classified FSMs of viruses into 2-indel compensatory FSMs and non-compensatory FSMs, and suggested that the frequencies of these two types of FSMs were similar (9, 24). Here we identified that, from a broader view, there are 3-, 4-, and 5-indel compensatory FSMs in virus evolution, besides 1-indel and 2-indel FSMs (Fig. 3A). We further showed that most FSMs in virus evolution are 2-indel compensatory FSMs and 3-indel compensatory FSMs are not rare (Fig. 3B).

This report demonstrated that FSMs usually occurred more frequently in virus genomes in the same family with lower sequence identities which could correspond to inter-species or inter-genus virus evolution. This finding suggests that FSMs are more important to the macroevolution or inter-species or inter-genus evolution of viruses than to the microevolution or intra-species evolution of viruses.

This report showed that FSMs occurred at distinct frequencies among virus families, which suggests that different virus families likely favor different mechanisms to change their amino acid sequences. For example, Table 1 suggests that paramyxoviruses likely employ FSMs to change their protein sequences more frequently than influenza viruses.

This report showed that FSMs occurred at distinct frequencies among different genes in the same virus genomes. This could result from the fact that different genes in the same viral genome undertake different positive selection pressure and negative selection pressure.

Fig. 2 showed that FSMs did not always occur more frequently in virus genomes of the same family with lower sequence identities. We think this could result from sampling errors, particularly when FSMs occurred at different frequencies among species in the family. Furthermore, FSM frequencies steeply increased when sequence identities declined to 60.0%−69.9% or 70.0%−79.9%, and the evolutionary significance of this phenomenon should be investigated in the future.

Because FSMs occurred frequently in the genomes of some species in some families, phylogenetic relationships of some viruses calculated with amino acid sequences could be somehow unreliable because one 2-indel compensatory FSM could be counted as multiple nucleotide substitutions through alignment of the relevant amino acid sequences (9).

Because FSMs can simultaneously change multiple amino acid residues, they can lead to major phenotypic transformations (i.e., macromutations), like virus genomic recombination, genomic reassortment, and mutations at key sites. These major phenotypic transformations could aid viruses to adapt to distinct environments or hosts, although most of them could reduce their fitness in their original environments or hosts. Therefore, this finding provided novel evidence to support the hopeful monster hypothesis in evolutionary biology, which claims that some major evolutionary transformations between species have occurred through large leaps of macromutations (24), rather than through the accumulation of small variations in Darwin’s theory.

The FSMs identified in this report were reliable only to a certain extent, because no alignment methods can fully and correctly reflect the real evolutionary history of nucleotide sequences. This issue is more obvious when sequence identities are lower (9). Therefore, we did not analyze FSM frequencies in the sequences with identities lower than 60.0% in this study. To improve the alignment quality and reduce the difficulty in analyzing alignment results, each fas file in this study contained only 2−10 sequences.

Table 3 suggests that the method using MAFFT with the mode of E-INS-I for identifying FSMs in this report was as stringent as the one using MUSCLE with the gap penalty of −800 and more stringent than other methods, which means that most FSMs identified in this report are reliable, although some FSMs could not have been identified. This does not affect the conclusions of this report because these conclusions were based on the same FSM identification method. Additionally, some FSMs identified in this study could result from sequencing errors. This problem usually cannot affect the conclusions of this report either, because these false FSMs should be distributed largely at random among the virus genes and families.

FSMs are widely distributed in the genomes of many organisms and can lead to phenotypic diversification (e.g., plumage variations) and pathological changes (e.g., cancers) in organisms (2, 3). However, few studies have been reported regarding the frequencies of FSMs in microevolution or macroevolution. Many viruses evolve much more rapidly than cellular organisms, and they are hence suitable to be employed to investigate various evolutionary theories and notions, including those regarding FSMs in microevolution or macroevolution. It is valuable to investigate in the future whether the features of FSMs in virus genomes revealed by this study are applicable in cellular organisms. This study could inspire researchers to investigate the roles, frequencies, and features of FSMs in the evolution of other viruses and cellular organisms, particularly on larger scales.

Together, this report described multiple fundamental features of FSMs in virus evolution for the first time. It also suggests that FSMs are important to the macroevolution and speciation of viruses, which is interesting in virology and evolutionary biology.

## MATERIALS AND METHODS

### Selection of viral families, genes, and sequences

Viral nucleotide sequences of complete ORFs of virus genes from 13 animal virus families were downloaded from National Library of Medicine (NCBI). The families and sequences were selected randomly. Some downloaded sequences were excluded due to poor sequencing quality (e.g., those sequences containing ambiguous nucleotides).

### Sequence alignment and sequence identity calculation

If not specified, nucleotide sequences were all aligned using the online software tool MAFFT (version 7) with the advanced setting of E-INS-I which is very slow and recommended for <200 sequences with multiple conserved domains and long gaps (25). Meanwhile, some fas files were aligned using the MUSCLE method incorporated in the software package MEGA (version 11) (26). Sequence identities were calculated using MEGA (version 11) as per pairwise deletion (26).

### Sequence grouping

Multiple fas sequence files were established for each virus family using the above randomly selected sequences. To improve the alignment quality and reduce the difficulty in parsing alignment results, each fas file contained only 2−10 sequences. Each fas files formed one or more sequence groups with intra-group sequence identities of 90.0%−99.9%, 80.0%−89.9%, 70.0%−79.9%, or 60.0%−69.9% (Fig. 1A) (Table S1). One sequence could be shared by two or more groups (Fig. 1A) (Table S1). Those sequences outside the above four sequence identity ranges were deleted. Some sequences were randomly added or deleted to make each of the 13 families have 10−50 groups for each of the four sequence identity ranges.

### FSM frequency calculation

When two or more adjacent frameshifting indels restored the original ORF, these adjacent indels constituted a compensatory FSM. Those FSMs constituted by only one indel were termed non-compensatory FSMs, which are usually near termination codons. FSM frequencies were calculated as per FSM counts, group counts, sequence average length, genes, and sequence identities. For example, if 90 sequences of a virus gene with 2700 nucleotides constituted 12 groups with intra-group sequence identities of 80.0%−89.9%, and these 12 groups contained 36 FSMs, then the FSM frequency of this gene on the scale of sequence identities of 80.0%−89.9% was 36/12/2,700*5,000 = 0.56 FSMs per group per 5,000 nucleotides. FSM starting and ending sites were numbered as per the reference sequences, which were randomly selected from the relevant groups (Table S1).

### Statistical tests

Data differences were compared using proper statistical tests (e.g., those groups of data that were not in the normal distribution were compared using the common or paired samples Wilcoxon tests). P<0.05 represented statistical significance. Results of statistical tests and the relevant original data were given in Text S1.

## ACKNOWLEDGMENTS

Many viral gene sequences in GenBank constituted the vital basis of this study. The authors thank deeply all the people who produced and shared these virus sequences in GenBank. This study was supported by the Open Competition Program of Top Ten Critical Priorities of Agricultural Science and Technology Innovation for the 14th Five-Year Plan of Guangdong Province (2022SDZG02). The funding agency did not play any role in this study.

## SUPPLEMENTARY MATERIAL

Supplemental material is available online only.

Supplemental file 1 Table S1 and Table S2

Supplemental file 2 Text S1

