## Supplementary material for "Frameshift Mutations in the Microevolution and Macroevolution of Viruses": Text S1

**Supplementary file**

**Text S1**

**Original data and relevant statistical tests**

**Table A1.** Frequencies of frameshift mutations (FSMs) in viral sequences with different sequence identities in 13 viral families identified in this study.

| **Virus family** | **FSM frequencies as per sequence identities** | | | |
| --- | --- | --- | --- | --- |
|  | **90.0%−99.9%** | **80.0%−89.9%** | **70.0%−79.9%** | **60.0%−69.9%** |
| *Arenaviridae* | 0.11 | 0.27 | 0.14 | 1.96 |
| *Circoviridae* | 0.00 | 0.00 | 2.66 | 3.58 |
| *Coronaviridae* | 0.18 | 0.39 | 0.91 | 3.58 |
| *Flaviviridae* | 0.07 | 0.16 | 0.10 | 0.89 |
| *Hantaviridae* | 0.09 | 0.15 | 0.22 | 1.27 |
| *Nairoviridae* | 0.00 | 0.00 | 0.45 | 1.88 |
| *Orthomyxoviridae* | 0.00 | 0.00 | 0.00 | 0.33 |
| *Paramyxoviridae* | 0.03 | 0.19 | 0.20 | 1.36 |
| *Peribunyaviridae* | 0.07 | 0.06 | 0.06 | 1.22 |
| *Phenuiviridae* | 0.00 | 0.00 | 0.00 | 1.20 |
| *Picornaviridae* | 0.07 | 0.30 | 0.61 | 1.94 |
| *Poxviridae* | 0.00 | 0.00 | 0.30 | 0.92 |
| *Togaviridae* | 0.07 | 0.06 | 1.53 | 2.85 |

**Table A2.** The Z-values and P-values of the paired samples Wilcoxon test used to compare the above four columns as per sequence identities.

| **Compared columns** | **Z-value** | **P-value** |
| --- | --- | --- |
| 90.0−99.9% versus 80.0−89.9% | -2.106 | 0.035 |
| 80.0−89.9% versus 70.0−79.9% | -2.191 | 0.028 |
| 70.0−79.9% versus 60.0−69.9% | -3.181 | 0.001 |

**Table B1.** Distribution of five types of FSMs identified in this study in 13 virus families

| **Virus family** | **Counts of different types of FSMs** | | | | |
| --- | --- | --- | --- | --- | --- |
|  | **1-indel** | **2-indels** | **3-indels** | **4-indels** | **5-indels** |
| *Arenaviridae* | 2 | 67 | 10 | 0 | 1 |
| *Circoviridae* | 0 | 11 | 0 | 0 | 0 |
| *Coronaviridae* | 1 | 66 | 3 | 0 | 0 |
| *Flaviviridae* | 0 | 44 | 5 | 0 | 0 |
| *Hantaviridae* | 0 | 29 | 1 | 1 | 0 |
| *Nairoviridae* | 0 | 86 | 6 | 2 | 0 |
| *Orthomyxoviridae* | 0 | 2 | 0 | 0 | 0 |
| *Paramyxoviridae* | 0 | 42 | 7 | 0 | 0 |
| *Peribunyaviridae* | 1 | 41 | 2 | 0 | 0 |
| *Phenuiviridae* | 0 | 14 | 1 | 0 | 0 |
| *Picornaviridae* | 0 | 117 | 8 | 1 | 0 |
| *Poxviridae* | 0 | 7 | 0 | 0 | 0 |
| *Togaviridae* | 0 | 81 | 19 | 1 | 0 |
| **Total** | **4** | **607** | **62** | **5** | **1** |

**Table B2.** The Z-values and P-values of the paired samples Wilcoxon test used to compare the above five columns as per FSM types.

| **Compared columns** | **Z-value** | **P-value** |
| --- | --- | --- |
| 1-indel versus 2-indel | -3.181 | 0.001 |
| 3-indel versus 2-indel | -3.181 | 0.001 |
| 4-indel versus 2-indel | -3.180 | 0.001 |
| 5-indel versus 2-indel | -3.181 | 0.001 |
| 1-indel versus 3-indel | -2.812 | 0.049 |
| 4-indel versus 3-indel | -2.668 | 0.008 |
| 5-indel versus 3-indel | -2.805 | 0.005 |
| 4-indel versus 1-indel | -0.264 | 0.792 |
| 5-indel versus 1-indel | -1.732 | 0.083 |
| 4-indel versus 5-indel | -1.414 | 0.157 |

**Table C1.** Counts of FSMs in viral sequences aligned using two software tools.

| **Virus genes** | **Sequences** | **Gap penalty_ MUSLE** | | |  | **MAFFT modes** | |
| --- | --- | --- | --- | --- | --- | --- | --- |
|  |  | **−400** | **−600** | **−800** |  | **E-INS-I** | **FFT-NS-2** |
| Spike protein of coronaviruses | NC004718  NC045512 | 3 | 0 | 0 |  | 0 | 1 |
| Large protein of nairoviruses | EU257628  NC004159 | 2 | 2 | 2 |  | 2 | 2 |
| Polyprotein of picornaviruses | AY843307  AY302559 | 8 | 1 | 1 |  | 1 | 1 |
| NS protein of togaviruses | NC001449  NC038670 | 5 | 2 | 2 |  | 2 | 2 |
| Large protein of paramyxoviruses | MH972568  NC006428 | 24 | 4 | 2 |  | 2 | 8 |
| Large protein of arenaviruses | NC010703  NC010702 | 10 | 2 | 1 |  | 1 | 2 |
| Polyprotein of flaviviruses | NC_043110  NC_033693 | 5 | 1 | 1 |  | 1 | 1 |
| Large protein of peribunyaviruses | KR149249  MG792213 | 23 | 7 | 2 |  | 2 | 9 |
| Large protein of hantaviruses | NC_005235  NC_005226 | 9 | 2 | 0 |  | 0 | 3 |
| Hemagglutinin of orthomyxoviruses | CY136872  EU030984 | 7 | 1 | 0 |  | 1 | 1 |
| Capsid of circoviruses | NC_055123  NC_026945 | 5 | 2 | 1 |  | 1 | 3 |
| Large protein of phenuviruses | NC027717  NC043611 | 13 | 4 | 0 |  | 1 | 4 |
| DNA polymerase of poxviruses | NC_031033  NC_035460 | 4 | 3 | 1 |  | 1 | 3 |

**Table C2.** The Z-values and P-values of the paired samples Wilcoxon test used to compare the above four columns as per alignment methods.

| **Compared columns** | **Z-value** | **P-value** |
| --- | --- | --- |
| −400 versus −600 (gap penalty) | -3.065 | 0.002 |
| −600 versus −800 (gap penalty) | -2.546 | 0.011 |
| −800 versus E-INS-I | -1.414 | 0.157 |
| E-INS-I versus FFT-NS -2 | -2.530 | 0.011 |

**Table D1.** FSM frequencies in the groups of four virus genes with sequence identities of 60.0−69.9% (their averages were given in Table 1). The frequencies were calculated using randomly selected sequences and statistically compared in the last line.

| **Gene** | Spike | Hemagglutinin | Polyprotein | Polyprotein |
| --- | --- | --- | --- | --- |
| **Family** | *Coronaviridae* | *Orthomyxoviridae* | *Flaviridae* | *Picornaviridae* |
| **Group number** | 20 | 18 | 21 | 33 |
| **FSM frequencies of each group** | 2.6; 1.3; 3.9; 3.9; 5.2; 1.3; 5.2; 5.2; 9.1; 5.2; 2.6; 1.3; 3.9; 6.5; 0.0; 3.9; 2.6; 2.6; 2.6; 2.6 | 0.00; 0.00; 0.00; 2.94; 0.00; 0.00; 0.00; 0.00; 0.00; 0.00; 0.00; 2.94; 0.00; 0.00; 0.00; 0.00; 0.00; 0.00 | 0.49; 1.48; 0.99; 0.49; 0.49; 0.49; 0.49; 0.00; 0.99; 2.47; 0.00; 2.47; 0.00; 0.00; 0.00; 0.49; 0.49; 0.99; 1.48; 1.98; 2.47 | 0.00; 2.23; 0.00; 1.49; 2.98; 0.74; 0.74; 3.72; 2.98; 1.49; 5.21; 1.49; 2.23; 4.46; 0.74; 1.49; 0.74; 2.23; 2.23; 2.23; 2.98; 2.98; 0.00; 0.00; 0.00; 1.49; 0.74; 0.00; 2.98; 6.70; 2.98; 2.98; 0.74 |
| **Statistics** | Z=-3.444, P=0.001  by the paired samples Wilcoxon test | | Z=-2.696, P=0.007  by the paired samples Wilcoxon test | |

**Table E.** FSM frequencies in a viral polymerase gene and a surface protein gene of 13 virus families with sequence identities of 60.0−79.9% (details of the FSMs in these genes are given in Table S2). The frequencies were calculated using randomly selected sequences. The frequencies were statistically compared in the last line.

| Family | Polymerase gene | Surface protein gene |
| --- | --- | --- |
| *Arenaviridae* | 1.7 | 13.9 |
| *Arenaviridae* | 2.3 | 0.0 |
| *Arenaviridae* | 1.5 | 0.0 |
| *Paramyxoviridae* | 1.5 | 6.1 |
| *Paramyxoviridae* | 3.0 | 3.0 |
| *Paramyxoviridae* | 0.7 | 2.5 |
| *Hantaviridae* | 0.8 | 7.3 |
| *Hantaviridae* | 2.3 | 10.2 |
| *Hantaviridae* | 0.8 | 7.3 |
| *Nairoviridae* | 0.8 | 8.0 |
| *Nairoviridae* | 1.7 | 6.0 |
| *Nairoviridae* | 0.4 | 7.6 |
| *Phenuviridae* | 1.6 | 5.1 |
| *Phenuviridae* | 0.8 | 0.0 |
| *Phenuviridae* | 2.4 | 9.7 |
| *Peribunyaviridae* | 0.7 | 0.0 |
| *Peribunyaviridae* | 0.7 | 9.4 |
| *Peribunyaviridae* | 0.8 | 5.9 |
| *Orthomyxoviridae* | 0.0 | 0.0 |
| *Orthomyxoviridae* | 0.0 | 2.9 |
| *Orthomyxoviridae* | 0.0 | 5.9 |
| *Circoviridae* | 0.0 | 6.9 |
| *Circoviridae* | 5.6 | 6.9 |
| *Circoviridae* | 0.0 | 7.2 |
| *Coronaviridae* | 0.0 | 4.0 |
| *Coronaviridae* | 1.1 | 2.5 |
| *Coronaviridae* | 0.8 | 3.8 |
| *Togaviridae* | 2.1 | 2.7 |
| *Togaviridae* | 2.0 | 5.4 |
| *Togaviridae* | 0.7 | 4.0 |
| *Poxviviridae* | 1.7 | 9.0 |
| *Poxviviridae* | 0.0 | 0.0 |
| *Poxviviridae* | 1.7 | 8.9 |
| *Picornaviridae* | 0.0 | 9.8 |
| *Picornaviridae* | 0.0 | 6.1 |
| *Picornaviridae* | 0.0 | 15.9 |
| *Flaviviridae* | 1.9 | 0.0 |
| *Flaviviridae* | 1.8 | 3.3 |
| *Flaviviridae* | 0.0 | 3.6 |
| Statistics | Z=-4.753, P=0.000  by the paired samples Wilcoxon test | |
